# Subject Instructions for Improved Characterization of Auditory Representations Using Reverse Correlation

**DOI:** 10.1101/2024.10.17.618874

**Authors:** Gidey W. Gezae, Nelson V. Barnett, Benjamin Parrell, Divya A. Chari, Adam C. Lammert

**Affiliations:** Human and Health Development, Pennsylvania State University, State College, PA, USA; Communication Sciences and Disorders, University of Wisconsin-Madison, Madison, WI, USA; Waisman Center, University of Wisconsin-Madison, Madison, WI, USA; University of Massachusetts Chan Medical School, Worcester, MA, USA; Massachusetts Eye and Ear Infirmary, Boston, MA, USA; Mathematics and Computer Science, College of the Holy Cross, Worcester, MA, USA; MGH Institute of Health Professions, Boston, MA, USA

## Abstract

Reverse Correlation (RC) is an established method for reconstructing auditory representations, that has recently emerged as a tool for characterizing the sounds experienced by tinnitus patients. Toward further optimizing RC for auditory research, the present work investigated the influence of subject instructions on characterization quality of tinnitus-like sounds. A validation study was conducted in which eighteen normal-hearing subjects were randomly assigned one of three candidate instruction sets, each inspired by the RC literature. Results show a significant effect of instruction set on characterization quality, and reveal that instructing subjects to detect a hidden signal in the RC stimuli resulted the best reconstruction.

## 1. Introduction

Internal auditory representations play an important role in mediating our perceptual experience of sound, by providing the critical link between the sensory signals available at the cochlea, and our perceptual experience of relatively abstract auditory objects. Auditory representations are used as the basis for interpreting sensory signals in accordance with various auditory percepts, ranging from low-level spatial and temporal features (e.g., pitches, timbres, formants), to higher-level cognitive constructs (e.g., phonemes, speaker identities, environmental sound sources). Auditory representations that are relatively localized in the auditory pathway (e.g., auditory receptive fields in primary auditory cortex) have seen substantial exposition, whereas auditory representations that are more distributed and/or abstract remain poorly understand by comparison, perhaps owing to a lack of approaches for measuring them directly.

One notable exception to the scarcity of approaches for directly measuring higher-level auditory representations is the method of Reverse Correlation (RC). RC asks subjects to render subjective judgments over random and/or ambiguous stimuli that have no direct relation to the representation of interest, from which unconstrained estimates of those representations can be rendered (Murray, 2011). While RC has seen applications throughout perceptual research, some of the earliest applications of RC were in auditory perception. Ahumada (1971) used RC to estimate perceptual representations of simple tones (Ahumada & Lovell, 2001). This pioneering work was later given a theoretical grounding based on a linear signal detection model by Richards & Zhu (1994). Auditory applications of RC have additionally gone beyond the perception of tones, to also look at the quality of more complex sounds, primarily in the context of speech perception. Brimijoin et al. (2013) utilized RC to perform unconstrained estimation of English vowels in 60 frequency bands spaced over the full audible range of 0.1-22kHz. Varnet and colleagues have applied RC in several studies aimed at uncovering the fine acoustic cues – due, e.g., to coarticulation – that promote robustness in speech recognition (Varnet, 2013a; Varnet 2013b; Varnet, 2015; Varnet, 2016). Ponsot (2018) has studied prosodic pitch variation associated with acoustic correlates of social judgments of, for example, dominance, trustworthiness, and smiling.

RC has also emerged recently as a promising method for characterizing tinnitus sounds. Tinnitus, a prevalent auditory condition characterized by the perception of sound without an external source, affects millions of people worldwide (Tunkel et al., 2014). Accurate characterization of tinnitus sounds across individuals is viewed as critical to progress in tinnitus research (Henry et al., 2013; Korth et al., 2021; Landgrebe et al., (2017); Lentz & He, 2020; Mohan et al., 2022, Ukaegbe et al., 2017), diagnosis (Chari & Limb, 2018; Heijneman et al., 2013; Norena et al., 2002; de Ridder et al., 2021; Roberts et al., 2008; Tyler et al., 1992), and treatment (Davis et al., 2007; Langguth et al., 2007; Nickel et al., 2005; Okamoto et al., 2010; Schaette et al., 2010; Stein et al., 2015; Tass et al., 2012). Recent evidence from applying RC to tinnitus has revealed that the method can characterize complex tinnitus sounds to a high level of accuracy (Hoyland, 2023; Barnett, 2024). Moreover, a fundamental concept underlying RC is that richly varying stimuli will, in some sense contain, information about the breadth of possible signals in the domain of interest – for auditory perception, the frequency domain. RC is thus especially appropriate for characterizing tinnitus given its wide heterogeneity in sound character (e.g,. roaring, buzzing, etc.), in contrast to existing methods – for example, pitch matching – that are best suited only for patients who experience tonal sounds (e.g., ringing) (Barnett, 2024).

However, it is not clear from these early applications whether RC protocols have been optimally adapted to the domain of tinnitus, specifically, or in the auditory domain, more broadly. RC protocols described in the research literature vary in several ways, including in the nature of the stimuli presented to subjects (e.g., Bernoulli-distributed (Gosselin & Schyns, 2003), or Gaussian-distributed (Ringach & Shapley, 2004), as well as pre-whitened (Compton, 2022) or templated (Dotsch & Todorov, 2012; Brinkman et al., 2017)), the method of estimation (e.g., linear (Gosselin & Schyns, 2003), sparse-constrained (Roop, 2021), or maximum *a posteriori* (Mineault, 2009)), and instructions given to subjects, the lattermost of which is the focus of the present study. While the subjective judgments that RC participants make are, ideally, well-informed by their internal perceptual experience, there are nevertheless many ways that participants might be instructed to listen, interpret, and ultimately respond to stimuli that are vague and ambiguous. The question of optimal subject instructions for RC seems especially pertinent in domains where the objects of interest are themselves often described as noisy – e.g., tinnitus sounds, or fricative sounds – which may confuse participants from the outset as to whether their task is one of finding *overt* matches to the percept of interest from among the stimuli, or whether their task is to identify aspects of similarity to the percept that may be *hidden* within the stimuli. Up to 50% of tinnitus patients may experience tinnitus sounds that are considered non-tonal, and may have noise-like qualities (Henry, 2013; Landgrebe, 2012; Lentz, 2020; Ukaegbe, 2017).

Overt matching asks subjects to identify noisy stimuli that simply match, to sufficient quality, the percept of interest. Perhaps owing to the fact that most applications of RC are not interested in percepts that are noisy in and of themselves, this approach is much less widely represented in the literature. A notable exception to this generalization, however, is represented by the overlapping literature on *pareidolia* (Hansen, 2010; Rekow, 2022; Salge, 2021), where the explicit aims is to study situations in which illusory percepts emerge from – as opposed to appearing “inside” or “underneath” – vague stimuli. Still, much more widely represented in the literature is the hidden approach to subject instructions, which typically involves asking subject to identify the percept of interest “within the noise.” Even within the hidden approach, there exist two broad subclasses of participant instructions in the literature: those based on *detection*, in which subjects identify stimuli containing the percept of interest from among those that contain no percept (Ahumada & Lovell, 1971; Gosselin & Schyns, 2003; Smith, 2012; Brimijoin, 2013), and those based on *discrimination*, in which subjects identify stimuli that contain the percept of interest from among that those contain other percepts (Brinkman, 2017; Ponsot, 2018; Varnet, 2022).

There is no reason to believe *a priori* that any of these approaches to RC subject instructions – overt, hidden/detection, or hidden/discrimination – will result in obviously more accurate characterization of auditory percepts, yet preliminary evidence from our research group, uncovered in the preparation of Hoyland et al. (2023) and unpublished, indicate that there may be reason to expect that subject instructions will have a measurable, and perhaps substantial, impact on the quality of the results. In any case, no direct comparison of subject instructions, and their effect on estimation quality, has been reported in the literature concerning RC for any sensory domain.

By investigating the impact of different instruction sets, we aim to identify the most effective instructions that enhance the validity and efficacy of RC for the reconstruction of auditory representations, in comprehending how sound characteristics contribute to tinnitus perception. Our approach involves the careful design of three distinct instruction sets in a validation study similar to that reported by Hoyland et al. (2003). The stimuli used were designed specifically to match potential qualities of tinnitus sounds, though the task and instructions are broadly applicable across the auditory domain. The results indicate that participant instructions have a large impact on the accuracy of RC reconstruction accuracy, with instructions consistent with a hidden/detection task providing the most accurate reconstructions.

## 2. Methods and Materials

Following the procedure described by Hoyland et al. (2023), we asked participants without tinnitus to complete an augmented RC experiment, in which they listened to both a target tinnitus-like sound in addition to a random-noise stimulus (described below) on every trial. Target sounds were subsequently used as the basis for validating the estimation quality of the RC-estimated tinnitus sounds by direct comparison.

### 2.1 Experimental Protocol

Subjects listened to A-X trials, containing a target sound (A) followed by a stimulus (X). X was randomly generated for each trial, while A remained the same across all trials (Fig. 1). Trials were presented in blocks of 100 trials, with voluntary breaks offered between blocks. Subjects completed five (5) blocks per target sound (p = 500 total trials per subject). Subjects listened over closed-back, circumaural earphones at a self-determined comfortable level in a quiet auditory environment. Presentation level was not recorded.

**Figure 1.**
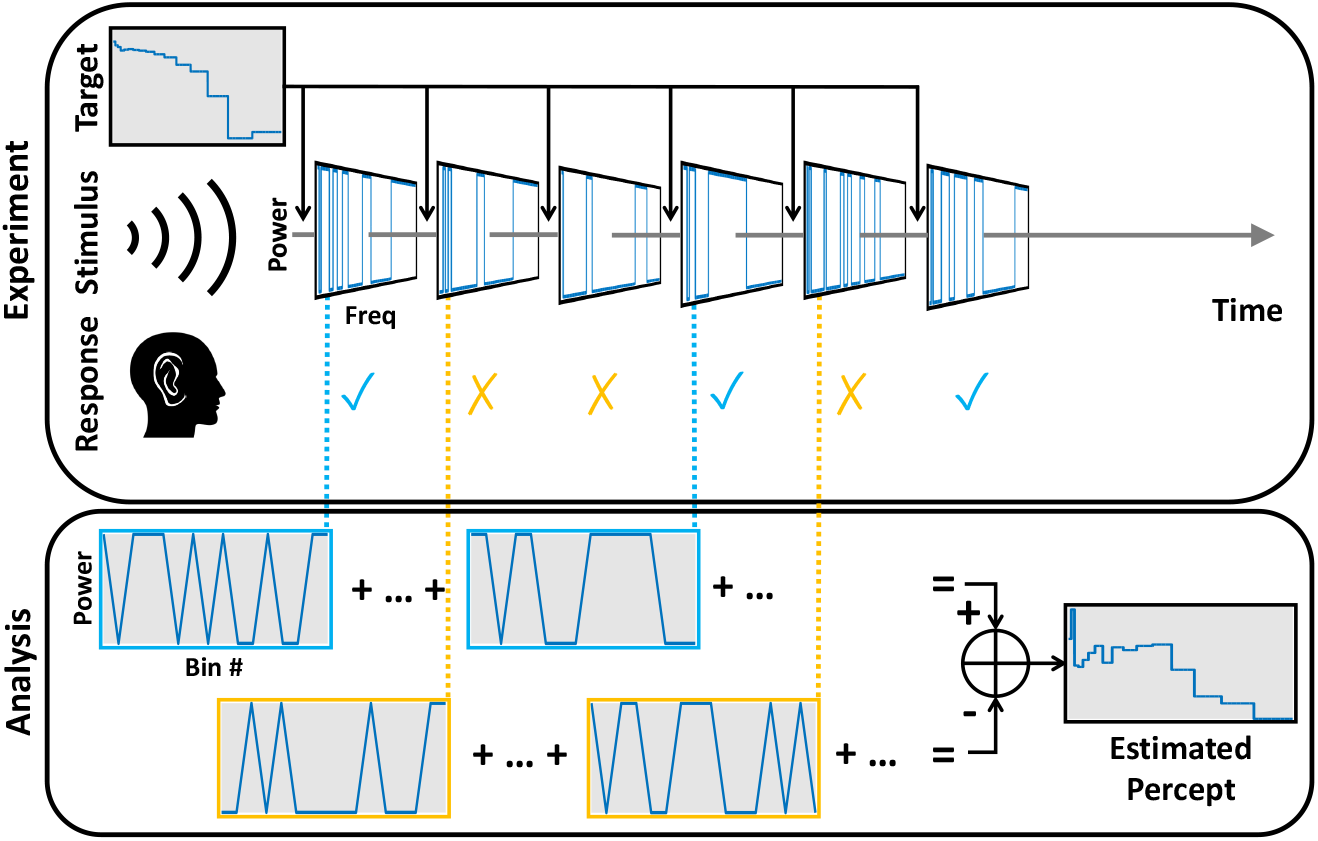
Illustration of the experimental protocol. Subjects listen to a sequence of randomly-generated auditory stimuli, each preceded by a target tinnitus-like sound. Subjects make a subjective judgment over the stimulus relative to the target sound, responding either “yes” or “no”, in light of the specific instruction set given to them. The resulting collection of all stimulus-response pairs is used to form an estimate of the target.

### 2.2 Participants

A total of 18 participants with no reported tinnitus or hearing loss were recruited for the study, ranging in age from 18 to 68 years. Participants were randomly assigned to one of three groups, with six participants in each group corresponding to the three different instruction sets described below. Procedures were approved by the UMass IRB.

### 2.3 Target Stimuli

Target sounds were drawn from online examples of tinnitus-like sounds maintained by the American Tinnitus Association (2022). The examples selected for the present study corresponded to the one described as “roaring.” This sound was selected for its spectra complexity, and due to its use in prior RC validation studies (Hoyland, 2023; Barnett, 2024). The sound was downloaded and truncated to 500 ms in duration. The purpose of these target sounds was primarily to serve as the representative tinnitus-like target for the purposes of estimation quality assessment for all participants, as detailed below.

### 2.4 Random-Noise Stimuli

Random-noise stimuli were generated as random frequency spectra, by dividing the range 100 Hz to 13,000 Hz into a total of 16 contiguous frequency bins with logarithmic spacing following the Mel scale. A minimum of 3 bins with a maximum of 9 bins were randomly set to 0dB amplitude and the rest were set to -100dB. Using these frequency spectra, each frequency was assigned random phase, and inverse Fourier transforms were subsequently used to compute audio waveforms of 500 ms duration corresponding to each spectrum.

### 2.5 Instruction Sets

In the first instruction set (Q1: hidden/detection), participants were tasked with listening to each random-noise stimulus, and providing a response based on the perceived presence or absence of the target “roaring” sound hidden in that stimulus. Subjects were told that a target sound would be hidden in approximately 50% the random-noise stimuli. A second instruction set (Q2: hidden/discrimination) asked subjects to listen to each random-noise stimulus, and provide a response based on whether the sound hidden in that stimulus was a good match to the target sound. Subjects were told that 100% of stimuli would contain a hidden sound, including a random assortment of roaring, buzzing, and ringing sounds. Finally, the third instruction set (Q3: overt) asked participants to listen to each random-noise stimulus, and provide a response based on whether the stimulus was a good match for the target sound. Note that none of the random-noise stimuli employed in the experiment contained any embedded sounds, nor were they intended to sound like the target, or indeed generated with any regard to the target sound whatsoever.

### 2.6 Tinnitus Sound Estimation

Estimates of individual participants percept of the target sound were obtained using the standard RC reconstruction procedure. That is, for each subject, given a 500-by-16 stimulus matrix X, where each row of X is a bin-power representation of one random-noise stimulus, and given a 500-by-1 response vector y, where entries in y have corresponding values of either -1 (“no” response) or +1 (“yes” response), estimates of the subject’s percept of the target sound may be calculated as β = 500^-1^X^T^y. It has been widely shown in the literature that this estimation formula is a restricted form of the Normal equation, which is the least-squares solution to linear regression (Gosselin & Schyns, 2003). In this specific case, the formula minimizes the sum of squared errors between the subject’s responses and the responses estimated using y_pred_ = X^T^β.

### 2.7 Estimation Quality Assessment

The power spectral density of the target sound was determined using Welch’s method (Welch, 1967), and subsequently transformed into a spectral representation allowing for direct comparison with the RC estimates. Frequency bins of the same number and spacing as those used for random-noise stimulus creation were defined, and the mean power at frequencies within each bin was bin-power representation, θ, that is calculated, resulting in a directly comparable to the bin-power representation of the random-noise stimuli. Quality of the estimates was quantified by correlating β with θ using Pearson’s *r*. P-values for each correlation were also calculated, and subsequently compared against the significance threshold of α = 0.05, corrected using the Holm-Bonferroni method. Differences in estimation quality between subjects given different instruction sets was tested using a one-way analysis of variance (ANOVA) to test for differences in the mean values of Fisher-tranformed (Fisher, 1915) Pearson’s r (see Table I) among the subjects groups receiving different instructions sets (i.e., Q1, Q2, or Q3). Further analysis using Tukey’s HSD (Honestly Significant Difference) Test was performed to compare the mean difference of estimation quality among the three instruction set groups.

**Table 1.** Pearson’s r and associated p values for the correlation between the RC estimate and the target sound spectrum for each participant. Bolded p-values indicate significant correlations at the Holm-Bonferroni correlated threshold value (α = 0.0028, DoF= 15).

| Instruction Set |  | Correlation Between Target and Reconstruction |  |  |  |  |  |
| --- | --- | --- | --- | --- | --- | --- | --- |
| Q1<br>(hidden/detect.) | Subject ID | S1 | S2 | S3 | S4 | S5 | S6 |
| | Pearson's $r$ | 0.819 | 0.786 | 0.706 | 0.535 | 0.495 | 0.344 |
| | $p$-value | **<0.001** | **0.001** | **0.003** | 0.040 | 0.061 | 0.210 |
| Q2<br>(hidden/discrim.) | Subject ID | S7 | S8 | S9 | S10 | S11 | S12 |
| | Pearson's $r$ | 0.635 | 0.615 | 0.327 | 0.292 | -0.403 | -0.588 |
| | $p$-value | 0.011 | 0.015 | 0.234 | 0.291 | 0.136 | 0.021 |
| Q3<br>(overt) | Subject ID | S13 | S14 | S15 | S16 | S17 | S18 |
| | Pearson's $r$ | 0.368 | 0.137 | -0.110 | -0.133 | -0.280 | -0.397 |
| | $p$-value | 0.177 | 0.626 | 0.696 | 0.637 | 0.312 | 0.143 |

## 3. Results

RC estimates of the participant’s percept of the target sound are shown in Figure 2, overlaid with a bin-power representation of the target sound spectrum. Qualitative differences in the correspondence between the estimates and the target sound can be seen across instruction set groups, as well as within those groups. The results of correlation analysis between each subject’s estimate and the target sound – including Pearson’s r and associated p-values – are listed in Table I. The only two subjects who showed statistically significant correlations were subjects S1 and S2, both of whom received instruction set Q1.

**Figure 2.**
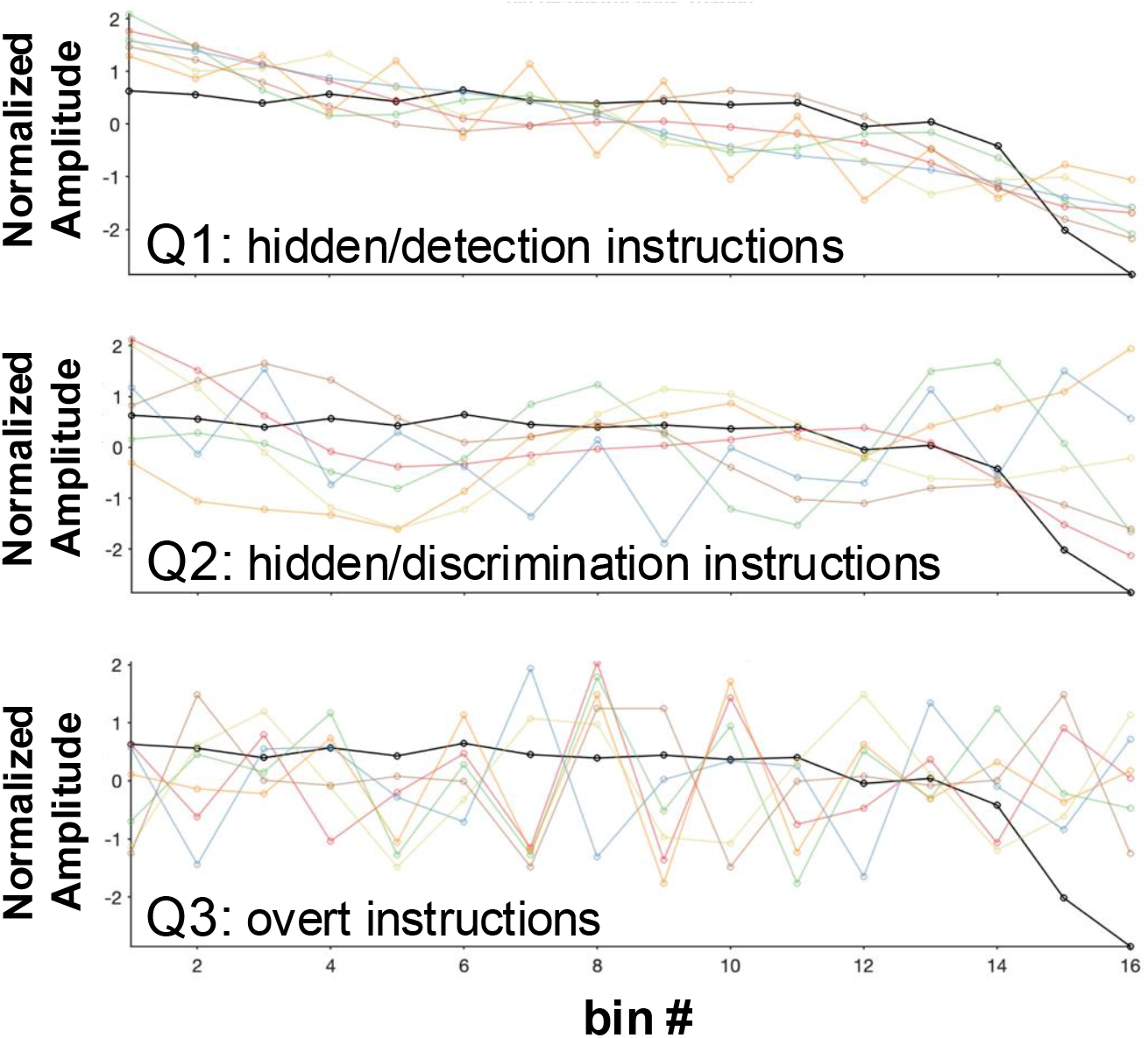
RC estimates of the target roaring sound from each of three instruction sets: Q1 (hidden/detection), Q2 (hidden/discrimination) and Q3 (overt). Subjects-specific estimates are indicated by the color traces. Black traces indicate the bin-power representation of the target sound. Power levels for each estimate were standardized prior to being plotted.

Participant instructions had a significant effect on the reconstruction accuracy, measured by the strength of the correlation between participants’ percepts and the true target spectra (F(2)=6.26, p=0.011, Figure 3). Post-hoc testing showed that Q1 (hidden/detection) led to significantly higher reconstruction accuracy than Q3 (overt; mean improvement = 0.147, q_s_(5)= 4.856, p = 0.010) There was no statistically significant difference between Q1 and Q2 (q_s_(5) = 3.470, p=0.066), or Q2 and Q3 (q_s_(5) = 1.386, p=0.600). In a follow-up analysis, it was found that the variance in reconstruction quality was significantly higher with Q2 (hidden/discrimination) than Q1 (hidden/detection) using a two-sample F-test for difference in variance between Q1 and Q2: F(5,5) = 0.1281, p = 0.042.

**Figure 3.**
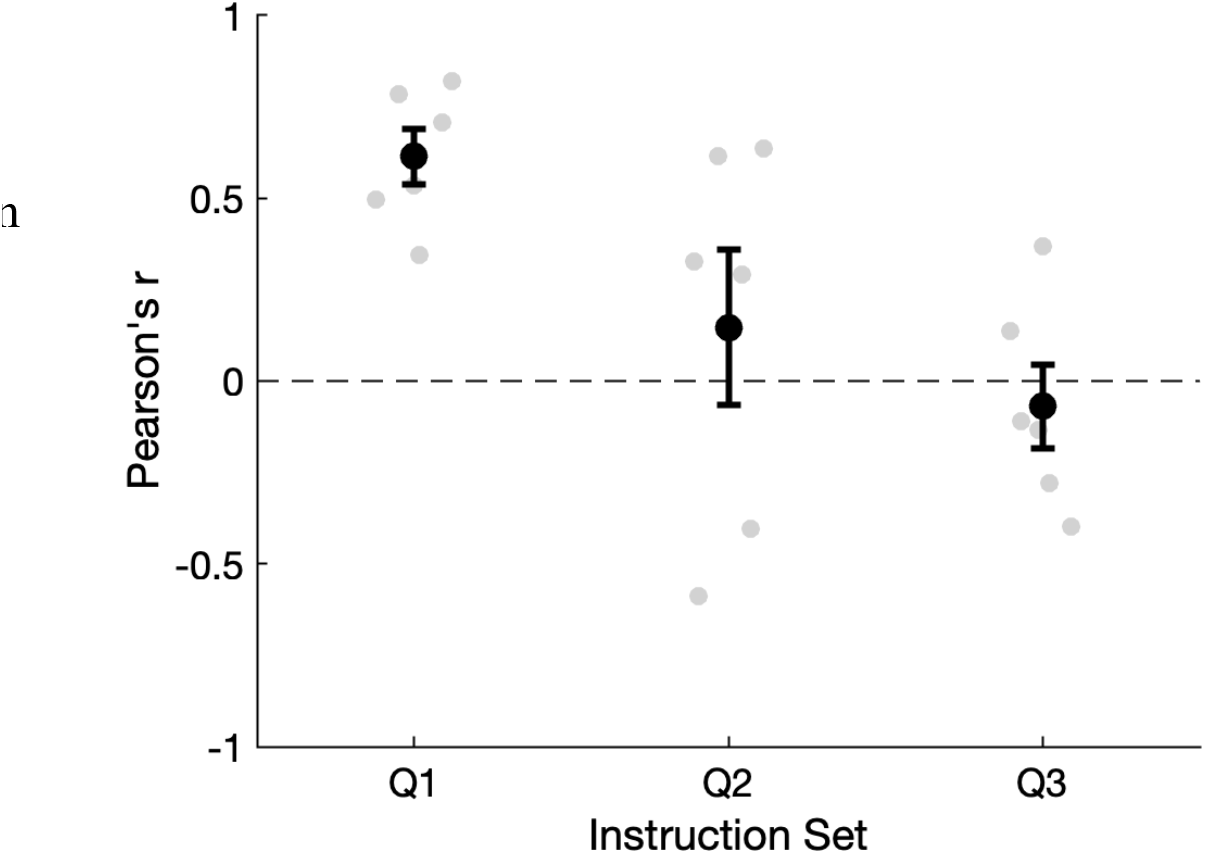
Scatter plot showing the distribution of Pearson’s r – a measure of RC estimation quality –for each instruction set as gray circles. Black circles indicate the mean value, and black lines indicate the upper and lower edge of the 95% confidence interval.

## 4. Discussion

We tested the effect of participant instructions on the accuracy of the reconstruction of auditory percepts in a signal detection task using reverse correction (RC). Our results show that instructions make a significant difference in the accuracy of these reconstructions, with the highest reconstruction accuracies coming from instructing subjects to detect the presence of the target sound hidden in a random-noise stimulus, which participants were told may or may not be present (Q1). Qualitatively, this approach resulted in better accuracy than instructions that told the participants to expect different embedded sounds on every trial, and subsequently respond when this embedded sound matched the target (Q2). Though this difference was not statistically significant, the variance in the accuracy of the reconstructions of the target sound was significantly higher with Q2 (hidden/discrimination) than Q1 (hidden/detection). The reason for this is unclear, but may be related to anecdotal reports from subjects that the Q2 instructions were confusing to understand. Finally, the hidden/detection instructions (Q1) provided a significant improvement over instructions to match the random noise stimulus and target sounds (Q3, overt).

While not the primary purpose of the study, it is also notable that these results represent the highest-resolution reconstructions of tinnitus-like sounds using RC reported in the literature, with the frequency resolution (i.e., the number of frequency bins in reconstructions) being double that reported by Hoyland et al. (2023). Moreover, these results are among the highest-reported reconstruction accuracies of tinnitus-like sounds, being comparable in estimation quality to lower-resolution reconstructions reported by Hoyland et al. (2023). That prior study also made use of Q1-type instructions, shown in this work to be the preferred set. Note that maintaining accuracy along with increased resolution is not necessarily expected, due to the widely observed tradeoff that exists in RC protocols between resolution and accuracy. Therefore, the comparable estimation quality observed in the present study are likely attributable to the 150% increase in the number of trials collected relative to Hoyland et al. (2023). Increasing the number of trials is a standard approach to accommodate the accuracy-resolution tradeoff.

The present study provides evidence for a preferred approach to instructing subjects engaged in auditory applications of Reverse Correlation, with regard to producing high quality reconstructions of auditory percepts. It was found that asking subject to detect the percept of interest embedded “within” the noisy stimuli, as distinguished from those stimuli where no such percept was embedded, provided estimates of the highest accuracy. This was shown in the context of reconstructing tinnitus-like sounds. However, it seems probable that subject instructions should be carefully considered in applications of RC to other aspects of auditory perception – e.g., the growing literature on RC in speech (Brimijoin, 2013; Varnet, 2016) – so as to preserve the language of the hidden approach used here. By refining the use of RC, researchers may be able to gain valuable scientific insights into how auditory phenomena are linked to personalized, cognitive representations of those phenomena. Improvements to RC may also enable improved diagnostic and treatment approaches for a variety of auditory conditions, by allowing more precise characterization of pathological representations (e.g., as in speech processing disorders), or by more effectively uncovering representations that are entirely internal to the individual (e.g., as in tinnitus).

## Acknowledgements

This work was supported by the University of Massachusetts Center for Clinical and Translational Science, and a grant from the American Tinnitus Association.

## References

Ahumada Jr, A., & Lovell, J. (1971). Stimulus features in signal detection. The Journal of the Acoustical Society of America, 49(6B), 1751–1756.

Ahumada, A. J. (2002). Classification image weights and internal noise level estimation. Journal of Vision, 2(1), 8–8.

Barnett, N. V., Hoyland, A., Chari, D. A., Parrell, B., & Lammert, A. C. (2024). Reverse Correlation Characterizes More Complete Tinnitus Spectra in Patients. IEEE Open Journal of Engineering in Medicine and Biology.

Brimijoin, O.W., Akeroyd, M. A., Tilbury, E., & Porr, B. (2013). The internal representation of vowel spectra investigated using behavioral response-triggered averaging. The Journal of the Acoustical Society of America, 133(2), EL118–EL122.

Brinkman, L., Todorov, A., & Dotsch, R. (2017). Visualising mental representations: A primer on noise-based reverse correlation in social psychology. European Review of Social Psychology, 28(1), 333–361.

Chari, D. A., & Limb, C. J. (2018). Tinnitus. Medical Clinics, 102(6), 1081–1093.

Compton, A., Roop, B. W., Parrell, B., & Lammert, A. C. (2022). Stimulus whitening improves the efficiency of reverse correlation. Behavior research methods, 1–9.

Davis, P. B., Paki, B., & Hanley, P. J. (2007). Neuromonics tinnitus treatment: third clinical trial. Ear and hearing, 28(2), 242–259.

Dotsch, R., & Todorov, A. (2012). Reverse correlating social face perception. Social Psychological and Personality Science, 3(5), 562–571.

Fisher, R. A. (1915). Frequency distribution of the values of the correlation coefficient in samples from an indefinitely large population. Biometrika, 10(4), 507–521.

Gosselin, F., & Schyns, P. G. (2003). Superstitious perceptions reveal properties of internal representations. Psychological science, 14(5), 505–509.

Hansen, B. C., Thompson, B., Hess, R. F., & Ellemberg, D. (2010). Extracting the internal representation of faces from human brain activity: An analogue to reverse correlation. NeuroImage, 51(1), 373–390.

Heijneman, K. M., De Kleine, E., Wiersinga-Post, E., & Van Dijk, P. (2013). Can the tinnitus spectrum identify tinnitus subgroups? Noise and Health, 15(63), 101.

Henry, J. A., Roberts, L. E., Ellingson, R. M., & Thielman, E. J. (2013). Computer-automated tinnitus assessment: noise-band matching, maskability, and residual inhibition. Journal of the American Academy of Audiology, 24(06), 486–504.

Hoyland, A., Barnett, N. V., Roop, B. W., Alexandrou, D., Caplan, M., Mills, J., … & Lammert, A. C. (2023). Reverse Correlation Uncovers More Complete Tinnitus Spectra. IEEE Open Journal of Engineering in Medicine and Biology.

Korth, D., Wollbrink, A., Lukas, C., Ivansic, D., Guntinas-Lichius, O., Salvari, V., … & Dobel, C. (2021). Comparing pure tone and narrow band noise to measure tonal tinnitus pitch-match frequency. Progress in Brain Research, 262, 115–137.

Landgrebe, M., Hajak, G., Wolf, S., Padberg, F., Klupp, P., Fallgatter, A. J., … & Langguth, B. (2017). 1-Hz rTMS in the treatment of tinnitus: A sham-controlled, randomized multicenter trial. Brain stimulation, 10(6), 1112–1120.

Langguth B, Goodey R, Azevedo A, et al. (2007). Consensus for tinnitus patient assessment and treatment outcome measurement: Tinnitus Research Initiative meeting, Regensburg, July 2006. Prog Brain Res 2007; 166: 525–36.

Lentz, J. J., & He, Y. (2020). Perceptual dimensions underlying tinnitus-like sounds. Journal of Speech, Language, and Hearing Research, 63(10), 3560–3566.

Listen to sample tinnitus sounds (ATA)”, Jun. 2022, [online] Available: https://www.ata.org/listen-sample-tinnitus-sounds.

Mineault, P. J., Barthelme, S., & Pack, C. C. (2009). Improved classification images with sparse priors in a smooth basis. Journal of Vision, 9(10), 17–17.

Mohan, A., Leong, S. L., De Ridder, D., & Vanneste, S. (2022). Symptom dimensions to address heterogeneity in tinnitus. Neuroscience & Biobehavioral Reviews, 134, 104542.

Murray, R. F. (2011). Classification images: A review. Journal of vision, 11(5), 2–2.

Nickel, A. K., Hillecke, T., Argstatter, H., & Bolay, H. V. (2005). Outcome Research in Music Therapy: A Step on the Long Road to an Evidence-Based Treatment. Annals of the New York Academy of Sciences, 1060(1), 283–293.

Norena, A., Micheyl, C., Chéry-Croze, S., & Collet, L. (2002). Psychoacoustic characterization of the tinnitus spectrum: implications for the underlying mechanisms of tinnitus. Audiology and Neurotology, 7(6), 358–369.

Okamoto, H., Stracke, H., Stoll, W., & Pantev, C. (2010). Listening to tailor-made notched music reduces tinnitus loudness and tinnitus-related auditory cortex activity. Proceedings of the National Academy of Sciences, 107(3), 1207–1210.

Ponsot, E., Burred, J. J., Belin, P., & Aucouturier, J. J. (2018). Cracking the social code of speech prosody using reverse correlation. Proceedings of the National Academy of Sciences, 115(15), 3972–3977.

de Ridder, D., Schlee, W., Vanneste, S., Londero, A., Weisz, N., Kleinjung, T., … & Langguth, B. (2021). Tinnitus and tinnitus disorder: Theoretical and operational definitions (an international multidisciplinary proposal). Progress in brain research, 260, 1–25.

Rekow, D., Baudouin, J. Y., Brochard, R., Rossion, B., & Leleu, A. (2022). Rapid neural categorization of facelike objects predicts the perceptual awareness of a face (face pareidolia). Cognition, 222, 105016.

Richards, V. M., & Zhu, S. (1994). Relative estimates of combination weights, decision criteria, and internal noise based on correlation coefficients. The Journal of the Acoustical Society of America, 95(1), 423–434.

Ringach, D., & Shapley, R. (2004). Reverse correlation in neurophysiology. Cognitive Science, 28(2), 147–166.

Roberts, L. E., Moffat, G., Baumann, M., Ward, L. M., & Bosnyak, D. J. (2008). Residual inhibition functions overlap tinnitus spectra and the rejavascript:void(0)gion of auditory threshold shift. Journal of the Association for Research in Otolaryngology, 9(4), 417–435.

Roop, B. W., Parrell, B., & Lammert, A. C. (2024). A compressive sensing approach for inferring cognitive representations with reverse correlation. Behavior Research Methods, 56(4), 3606–3618.

Salge, J. H., Pollmann, S., & Reeder, R. R. (2021). Anomalous visual experience is linked to perceptual uncertainty and visual imagery vividness. Psychological research, 85(5), 1848–1865.

Schaette, R., König, O., Hornig, D., Gross, M., & Kempter, R. (2010). Acoustic stimulation treatments against tinnitus could be most effective when tinnitus pitch is within the stimulated frequency range. Hearing Research, 269(1-2), 95–101.

Stein, A., Engell, A., Junghoefer, M., Wunderlich, R., Lau, P., Wollbrink, A., … & Pantev, C. (2015). Inhibition-induced plasticity in tinnitus patients after repetitive exposure to tailor-made notched music. Clinical Neurophysiology, 126(5), 1007–1015.

Tass, P. A., Adamchic, I., Freund, H. J., von Stackelberg, T., & Hauptmann, C. (2012). Counteracting tinnitus by acoustic coordinated reset neuromodulation. Restorative neurology and neuroscience, 30(2), 137–159.

Tunkel, D. E., Bauer, C. A., Sun, G. H., Rosenfeld, R. M., Chandrasekhar, S. S., Cunningham Jr, E. R., … & Whamond, E. J. (2014). Clinical practice guideline: tinnitus. Otolaryngology– Head and Neck Surgery, 151(2_suppl), S1–S40.

Tyler, R. S., Aran, J. M., & Dauman, R. (1992). Recent advances in tinnitus. American journal of audiology, 1(4), 36–44.

Ukaegbe, O. C., Orji, F. T., Ezeanolue, B. C., Akpeh, J. O., & Okorafor, I. A. (2017). Tinnitus and its effect on the quality of life of sufferers: a Nigerian cohort study. Otolaryngology– Head and Neck Surgery, 157(4), 690–695.

Varnet, L., Knoblauch, K., Meunier, F., & Hoen, M. (2013a). Using auditory classification images for the identification of fine acoustic cues used in speech perception. Frontiers in human neuroscience, 7, 865.

Varnet, L., Knoblauch, K., Meunier, F., & Hoen, M. (2013b). Show me what you listen to! Auditory classification images can reveal the processing of fine acoustic cues during speech categorization. In INTERSPEECH (pp. 3167–3171).

Varnet, L., Knoblauch, K., Serniclaes, W., Meunier, F., & Hoen, M. (2015). A psychophysical imaging method evidencing auditory cue extraction during speech perception: a group analysis of auditory classification images. PLoS One, 10(3), e0118009.

Varnet, L., Meunier, F., Trollé, G., & Hoen, M. (2016). Direct viewing of dyslexics’ compensatory strategies in speech in noise using auditory classification images. PloS one, 11(4), e0153781.

Welch, P. (1967). The use of fast Fourier transform for the estimation of power spectra: a method based on time averaging over short, modified periodograms. IEEE Transactions on audio and electroacoustics, 15(2), 70–73.

